# Stratification of amyotrophic lateral sclerosis patients: a crowdsourcing approach

**DOI:** 10.1101/294231

**Authors:** Robert Kueffner, Neta Zach, Maya Bronfeld, Raquel Norel, Nazem Atassi, Venkat Balagurusamy, Barbara di Camillo, Adriano Chio, Merit Cudkowicz, Donna Dillenberger, Javier Garcia-Garcia, Orla Hardiman, Bruce Hoff, Joshua Knight, Melanie L. Leitner, Guang Li, Lara Mangravite, Thea Norman, Liuxia Wang, the ALS Stratification Consortium, Jinfeng Xiao, Wen-Chieh Fang, Jian Peng, Gustavo Stolovitzky

**Affiliations:** Icahn School of Medicine at Mount Sinai, New York, NY, USA; Teva Pharmaceuticals, Netanyah, Israel; Prize4Life, Haifa, Israel; IBM Research, Yorktown Heights, NY, USA; Massachusetts General Hospital, Boston, MA, USA; University of Padova, Padova, Italy; ‘Rita Levi Montalcini’ Department of Neuroscience, University of Turin, Turin, Italy, and Institute of Cognitive Sciences and Technologies, C.N.R, Rome, Italy; Pompeu Fabra University, Barcelona, Spain; Trinity College Institute of Neuroscience, Dublin, Ireland; Sage Bionetworks, Seattle, Washington, USA; Accelerating NeuroVentures, Boston, MA, USA; Amazon, Seattle, Washington, USA; Zillow, Seattle, Washington, USA; Department of Computer Science, University of Illinois at Urbana-Champaign, USA; Department of Information and Learning Technology, National University of Tainan, Tainan City, Taiwan; Department of Human Genetics, McGill University, Montreal, Canada; Ontario Institute for Cancer Research (OICR), Toronto, Canada; Departement d’Economie Ecole Polytechnique, Paris, France; Interdisciplinary Computing and Complex BioSystems (ICOS) research group, Newcastle University, UK; Veristat Inc, Southborough, MA; Medical Research, Kfar Malal, Israel; Department of Computer Science, Ben-Gurion University of the Negev, Israel; Department of Industrial Engineering and Management, Ben-Gurion University of the Negev, Israel; Faculty of Information Technology, Monash University, Australia; Analytica Laboratories, Hamilton, New Zealand; Department of Computer Science and Information Engineering, National Cheng Kung University, Taiwan; Princess Margaret Cancer Centre, Toronto, Ontario, Canada; Department of Biostatistics, University of Kansas Medical Center, Kansas City, KS; MIT, Department of Mathematics, Cambridge, MA; Perelman School of Medicine, University of Pennsylvania, Philadelphia, PA; Centre de Recherche en Economie et Statistique (CREST), Paris, France; Department of Biostatistics and Bioinformatics, Moffitt Cancer Center, Tampa, FL; Program on Bioinformatics and Systems Biology, Sanford Burnham Prebys Medical Discovery Institute, La Jolla, CA; Institute of Informatics, University of Bialystok, Bialystok, Poland; Department of Engineering, University of Cambridge, UK; RTI International, Research Triangle Park, NC; KU Leuven, Department of Public Health and Primary Care, Kortrijk, Belgium; Origent Data Sciences, Inc., Vienna, VA; National Center for Mathematics and Interdisciplinary Sciences, Academy of Mathematics and Systems Science, Chinese Academy of Sciences, China; Stanford University, Center for Biomedical Informatics Research, Stanford, CA; Department of Statistics, Tel-Aviv University, Israel; Helmholtz Zentrum Munchen, Institute of Health Economics and Health Care Management, Munich, Germany; Department of Bioinformatics, University of Bialystok, Ciolkowskiego, Bialystok, Poland; Shanghai Key Lab of Intelligent Information Processing and School of Computer Science, Fudan University, Shanghai, China; School of Computer, Wuhan University, Wuhan, China; Computational Centre, University of Bialystok, Ciolkowskiego, Bialystok, Poland; Children’s Mercy Hospital, Kansas City, MO; LinkedIn, Sunnyville, CA; Department of Computer Science, University of Illinois at Urbana-Champaign, IL; Interdisciplinary Centre for Mathematical and Computational Modelling, University of Warsaw, Pawinskiego, Warsaw, Poland; Department of Biomedical Engineering, School of Advanced Technologies in Medicine, Isfahan University of Medical Sciences, Isfahan, Iran; Ceremade Universite Paris-Dauphine, France, Université Paris-Est, Laboratoire d’Analyse et de Mathématiques Appliquées, Créteil, France; Stanford University, Department of Statistics, Stanford, CA; Stanford University, Department of Electrical Engineering, Stanford, CA; Department of Computer Science and Engineering, University of Moratuwa, Sri Lanka; Department of Computer Science, University of California, Irvine, CA; Paul G. Allen School of Computer Science & Engineering, University of Washington, Seattle, WA; Department of Computer Science, City University of Hong Kong, China; Dept of Computer Science, Bren School of Information and Computer Sciences, University of California, Irvine, CA; University of Pennsylvania, Philadelphia, PA; University of Maryland, Baltimore, MD; Centre for Computational System Biology, ISTBI, Fudan University, Shanghai, China

## Abstract

Amyotrophic lateral sclerosis (ALS) is a fatal neurodegenerative disease with substantial heterogeneity in clinical presentation with an urgent need for better stratification tools for clinical development and care. In this study we used a crowdsourcing approach to address the problem of ALS patient stratification. The DREAM Prize4Life ALS Stratification Challenge was a crowdsourcing initiative using data from >10,000 patients from completed ALS clinical trials and 1479 patients from community-based patient registers. Challenge participants used machine learning and clustering techniques to predict ALS progression and survival. By developing new approaches, the best performing teams were able to predict disease outcomes better than currently available methods. At the same time, the integration of clustering components across methods led to the emergence of distinct consensus clusters, separating patients into four consistent groups, each with its unique predictors for classification. This analysis reveals for the first time the potential of a crowdsourcing approach to uncover covert patient sub-populations, and to accelerate disease understanding and therapeutic development.

---

Amyotrophic lateral sclerosis (ALS) is a neurodegenerative disorder which causes the death of motor neurons that control voluntary muscles. The loss of motor neurons leads to progressive muscle weakening and paralysis and on average patients will survive only 3-5 years from symptom onset^1^. Despite being known for over 150 years, we only have limited understanding of the biological mechanisms underlying ALS and existing therapeutic options merely extend survival by a few months^2,3^. One of the biggest challenges in ALS treatment and research today is the well-established heterogeneity of the disease^4,1^; ALS patients can have widely different patterns of disease manifestation and progression, and genetic analyses suggest heterogeneity of the underlying biological mechanisms as well^5,6,7,8^. This heterogeneity has detrimental effects on clinical trial planning and interpretation, as it might mask drug effects^3^, on attempts to uncover disease mechanisms, and on clinical care, as it increases uncertainty about prognosis and makes treatment course planning challenging. Thus, successfully stratifying ALS patients into clinically meaningful sub-groups can be of great value for advancing the development of effective treatments and achieving better care for ALS patients.

Early classification systems for ALS patients were based on clinical presentation of the disease and were intended for ascertainment of an ALS diagnosis, but were limited in their ability to predict disease prognosis or suggest underlying disease mechanisms^9,10,4^. More recent attempts of ALS patient classification focused on prediction of clinical outcomes but were often limited by small sample sizes lacking the highly needed detailed characterization of patient subgroups^11,12,13,14^. In the current study, we sought to use the power of state of the art machine learning algorithms applied to a large-scale, clinically detailed database of ALS patients to uncover and characterize clinically significant subpopulations of ALS patients.

To address the need for a large, diverse and clinically relevant dataset, we used two complementary data sources. The first was data from ALS national or regional registers from Ireland and the Piemonte and Valle d’Aosta region in Italy, collected as part of standard clinical visits and representing ALS community data. The second dataset was ALS clinical trial data, from the Pooled Resource Open-Access ALS Clinical Trials platform (PRO-ACT, www.ALSdatabase.org/), an open-access database containing harmonized and de-identified data of over 10,000 ALS patients from 23 completed clinical trials^15^.

The PRO-ACT database was established in collaboration between Prize4Life (Haifa, Israel) and the Neurological Clinical Research Institute (NCRI) at Massachusetts General Hospital (Boston, MA, USA) with the purpose of advancing ALS research through ‘big data’ initiatives. The PRO-ACT database was previously used for a crowdsourcing computational challenge: The 2012 DREAM-Phil Bowen ALS Prediction Prize4Life challenge (The ALS Prediction Challenge)^16^. The ALS Prediction Challenge invited participants to develop computational algorithms that could predict ALS disease progression based on data collected during the first three months of clinical observation. The best performing algorithms developed for the challenge were able to achieve a prediction accuracy that would allow a 20% reduction in the number of patients needed for a trial^16^ and are currently being tested for use in clinical trials^17,18^.

Building upon the success of the earlier prediction challenge, the 2015 DREAM ALS Stratification Prize4Life Challenge (The ALS Stratification Challenge) was a crowdsourcing initiative that sought to extend the scope of prediction algorithms by inviting participants to stratify the ALS patient population into distinct clusters and develop separate predictive models for each subpopulation. The ALS Stratification Challenge included both disease progression and survival as predicted outcome measures and using either clinical trial or community registries data. This was designed to increase the relevance and applicability of predictive models to a broader patient population, to flush out any differences between the general patient population and the clinical trial population and to uncover clinical markers with high predictive value which are not collected as part of current routine clinical practice. Thus, the challenge was divided into four sub-challenges in which participants predicted either disease progression or survival while using data from either PRO-ACT or ALS registries. A prize of $28,000, collected through a crowdfunding effort, was divided equally between best performers of the four sub-challenges.

In this publication we describe the results of the challenge including analysis of the best performing algorithms’ performance and methods as well as findings regarding patient sub-populations and relative importance of different predictive features obtained from cross-model assessment.

## Results

### Challenge design

The ALS Stratification Challenge was developed and ran through a collaboration between the nonprofit organizations Dialogue for Reverse Engineering Assessments and Methods initiative (DREAM, http://dreamchallenges.org/) and Prize4Life (www.prize4life.org.il) using the Sage Bionetworks Synapse platform (www.synapse.org). The challenge was based on two datasets: (1) ALS clinical trials data collected through the PRO-ACT database, and (2) community-based ALS clinical data collected through ALS registries. Both datasets contained longitudinally sampled demographic and clinical information with some additional genetic (specific mutation) and family history data in the registries and detailed laboratory tests in PRO-ACT (See Supplementary material part 1 and 2 for detailed description of both datasets and the challenge description as given to participants, respectively).

The challenge was divided into four sub-challenges: (1) Predicting disease progression or (2) survival probability using PRO-ACT data and (3) predicting disease progression or (4) survival probability using ALS registries data. Challenge participants were asked to use patients’ data from the first 3 months of records to predict disease progression at 12 months or probability of survival at 12, 18 & 24 months. Disease progression was defined as the slope of the ALS Functional Rating Scale (ALSFRS or ALSFRS-R) between 3 and 12 months (see online methods). The challenge ran between June and October 2015 and drew 288 registrants, eventually leading to final submissions by 30 teams (88 individual participants) from 15 countries (see Supplementary material part 3 for participant survey).

### Approach to performance assessment

To avoid overfitting of algorithms^19^, two datasets were used for each sub-challenge, a training dataset accessible to participants in full, and a validation set, which was only available to the challenge organizers and was used to evaluate the prediction accuracy of submitted algorithms at the end of the challenge. For the PRO-ACT sub-challenges, data from the publicly available PRO-ACT database was used as the training set, and a separate set of data from six additional clinical trials which were not previously publicly available, was used for validation. The registry data, which was never before made publicly available, was divided randomly (split evenly across the two registries) into training and validation sets.

The accuracy of disease progression predictions was assessed by the combination of three different metrics: concordance index (CI), Pearson’s correlation coefficient (PCC) and root mean squared deviation (RMSD). The evaluation of survival prediction accuracy was based on the CI scores for the three survival probability predictions (probability of survival at 12, 18 & 24 months) and confirmed by a time-ROC analysis. The three scores for each sub-challenge were transformed into z-scores that were then combined to obtain a ranking of the submitted algorithms (see online methods). Performance was also compared to two baseline algorithms which were based on the top performing prediction algorithms submitted to the 2012 ALS prediction challenge^16^, adapted to the requirements of the new challenge (see Supplementary material part 4).

Clustering patients into subgroups was not mandated, but participants were encouraged to use clustering to enhance performance. The challenge introduced an additional requirement that predictions are to be made based on a limited number of clinical features. The requirement for limiting the number of features was highlighted by our clinical advisors to facilitate the application of predictive algorithms in natural clinical setting^13^. A preliminary analysis indicated the benefit from clustering (in terms of improved prediction accuracy) tends to increase when the number of features is restricted, and this effect plateaued at around 6 features (see Supplementary material part 4). Thus, participants were asked to write algorithms that first selected the most informative features (up to 6 features), and in a second step used only the data from these selected features to make progression or survival predictions (Figure 1). To enforce the separation between these two steps as well as to completely hide the validation set we required participants to implement their algorithms in Docker containers (https://docs.docker.com/) executed on the IBM Z cloud (https://www.ibm.com/it-infrastructure/z/capabilities/enterprise-security). This way, we could run the algorithms in a secured environment where the participants could not see the validation data or other jobs running.

**Figure 1:**
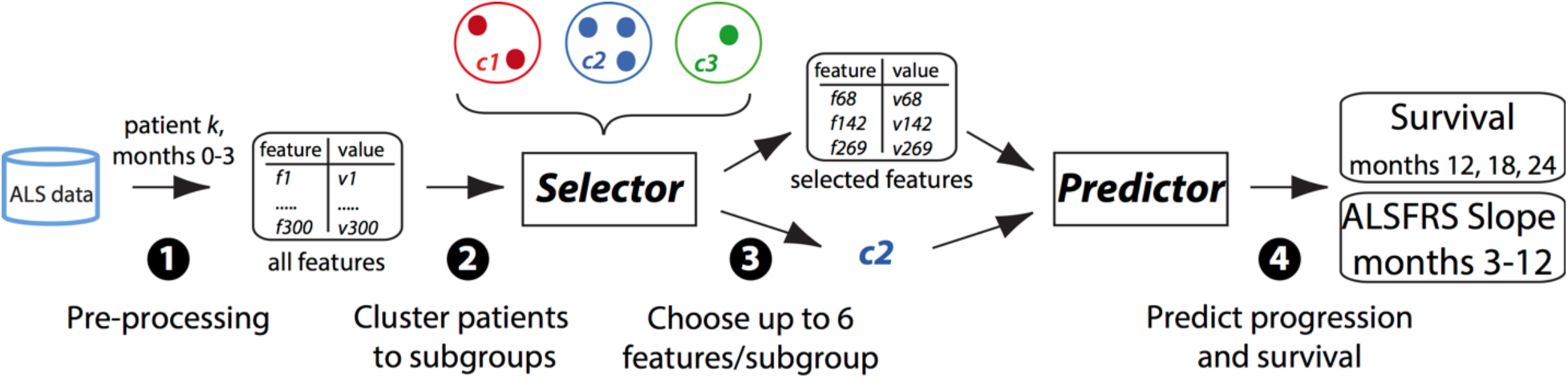
Outline of algorithms design. Algorithms used either PRO-ACT or ALS registries data, and first (1) applied various data pre-processing methods, such as data imputation, aggregation, quantization and more. Next, (2) algorithms could cluster the patient population into any number of sub-groups and (3) select the most informative features for each cluster (up to a maximum of 6 features). Then (4) a “predictor” component had to use values of the selected features to predict either disease progression or survival for any given patient. In the scoring of the challenge, the algorithms made predictions for patients that were not part of the original datasets available for algorithms training, and the accuracy of these predictions was assessed.

### Comparative assessment of prediction methods

In general, the top performing teams in each sub-challenge (except for the registry progression sub-challenge) significantly outperformed the best baseline algorithm (Figure 2, See Supplementary material part 5 for full results). Random forest was by far the most commonly used prediction method, with overall very good results. It was the method used by the best performing team in the registry progression sub-challenge and by most algorithms ranked between the 2nd and 8th places across all four sub-challenges (Figure 2a). Another successful method included a Gaussian process regression model with an arithmetic mean kernel, which was the best performing for the two survival challenges but did not perform so well for the progression challenges.

**Figure 2:**
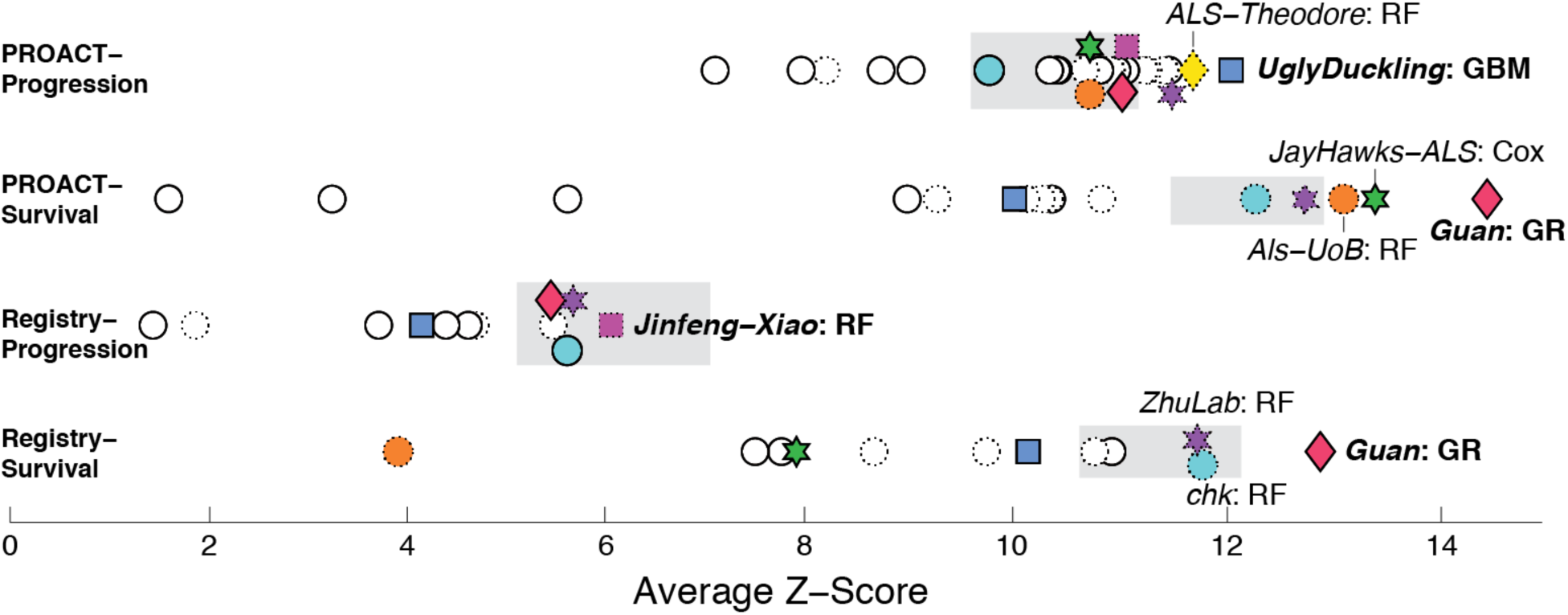
Overview of the performance of submitted and baseline algorithms across the four sub-challenges. Submissions were assessed by Z-scores combining RMSD, concordance index and Pearson’s correlation for the progression sub-challenges and three concordance indices for the survival sub-challenges. Grey boxes denote the performance of the best-performing baseline algorithm (left and right boundary of the box represent intervals of its performance ± the bootstrapped standard deviation). Teams that achieved the top three scores in any sub-challenge are indicated by colored symbols and shown by the same symbol in all sub-challenges. The underlying method is indicated (RF=random forest, GBM=generalized boosting model, Cox=Cox model, GR=Gaussian regression). Submissions based on random forests, the most frequently used method, are denoted by symbols with dashed outlines.

The performance of the top teams, as was observed in other crowdsourcing challenges^20,21^, varied substantially even if the same machine learning method was used, and depended to a large extend on data pre-processing, especially feature selection and representation of time-resolved features. Here, the best performing teams followed the successful logic of the previous ALS prediction prize^16^, where time-resolved information was represented by a combination of simple summary statistics (for instance minimum, maximum and average of the feature values). Due to the restriction of 6 features used for the prediction of a patient’s outcome, teams also needed to devise a suitable feature selection strategy. This was achieved by evaluating the contribution of the complete set of selected features (as opposed to selecting them one by one), for example through evaluating sets of features by their combined information gain or by aggregating their weight along the paths of all trees in a random forest.

Survival predictions deserve special consideration, as one team (same in both survival sub-challenges) out-performed other participants substantially. Survival predictions are particularly challenging due to the right-censored outcome variable survival time: data can be terminated by either patient death or by trial drop out. The standard Cox proportional hazards model, routinely used to explore the dependency between clinical features and survival, ignores such censored cases, which likely deteriorates performance. In the current challenge, Yuanfang Guan circumvented this problem via a novel strategy of incorporating Kaplan-Meier curves into Gaussian regression models. This defined the outcome variable more precisely, which led to a more adequate training of regression models (here: Gaussian process models) explaining the algorithm’s superior prediction performance. In line with these results, the strategy outperformed a standard Cox model by 20% accuracy^22^.

Notably, for the sub-challenges running on the PRO-ACT database, the best performing algorithms significantly outperformed the winning method from the first challenge^16^. This is even more noteworthy given that the validation set was not randomly divided from the training set, but actually included the more difficult and realistic criteria of application of the algorithms to a data comprising of six new never used before trials.

#### Predictive clinical features

The challenge’s requirement of using only up to six features for prediction encouraged participants to identify the most informative features for the prediction of disease progression or survival. We assessed the number of times each feature was used for prediction across all submitted algorithms and identified the features most frequently used within and across the different sub-challenges (Figure 3).

**Figure 3:**
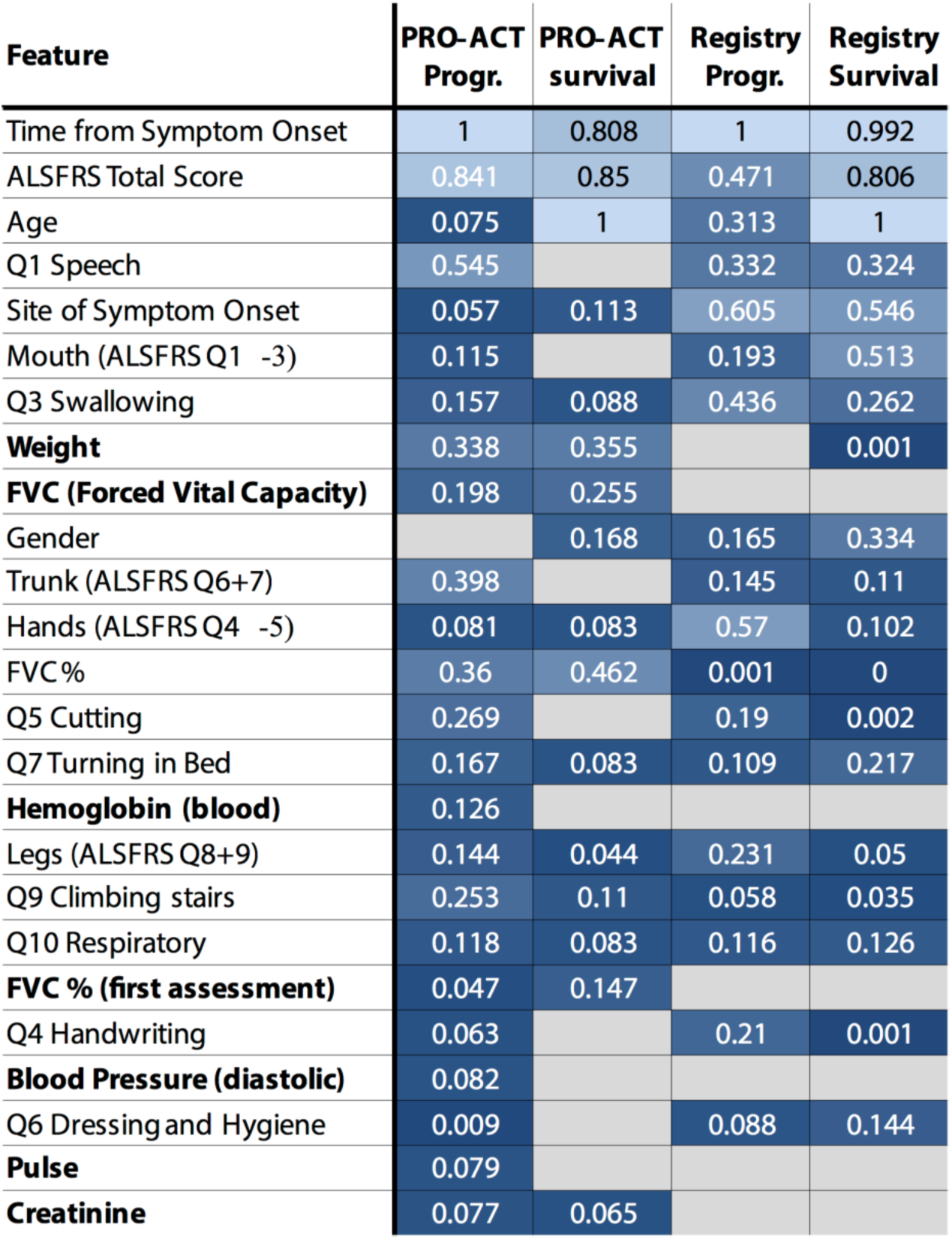
Overview of the features used by the algorithms. For each subchallenge, the listed probability shows how often the corresponding feature is used by the algorithms. The features are ranked-ordered by this probability, averaged across all sub-challenges where darker colors denote lower probabilities). Cases where a given feature was not used at all for a given sub-challenge are shown in grey (probability 0). Features that are recommended to be assessed by clinicians more often to aid prognosis are marked in bold.

The most frequently used features for all sub-challenges were those that are well described in the literature as being strongly related to ALS prognosis: time from disease onset and total ALSFRS score^23,24,25^. Age and gender were also frequently used but were more informative for predicting survival rather than disease progression, in line with literature^22, 23,24, 26,27,28^. While age and gender were generally predictive in this data as well, they were not specifically more predictive for one particular subgroup of patients, and therefore less relevant for stratification, while still not deterring from its overall importance as a predictor. Bulbar function is also known to be particularly informative for clinical outcome prediction^24,25,26^ and ALSFRS questions 1 and 3 (bulbar functions) were selected more frequently compared to any other functional domains across all sub-challenges. Interestingly, ALSFRS question Q2 (salivation) was rarely used, in line with previous works indicating this question might be less well correlated with total ALSFRS scores and/or disease progression, due to effective available treatment options^29^.

Data availability was another important factor for feature selection. For example, while weight or BMI, is known to correlate with ALS prognosis^30,31^ and was frequently selected for predictions for the PRO-ACT dataset, it was recorded in only <10% of the cases in the registry data, making it unusable for predictions (Fig. 3). The potential predictive benefit of features not routinely collected in clinical practice was also observed with respect to features evaluating breathing capacity. Given that loss of breathing function is the main cause of death for ALS patients^32^, it is not surprising that breathing-related features, primarily Forced Vital Capacity (FVC) were commonly used for predictions in both PRO-ACT sub-challenges. However, in the registry data, where FVC data was not available and breathing was only assessed by the ALSFRS scale breathing-related questions were less frequently used for prediction. This suggests that clinicians could potentially gain better insight into individual patient prognosis by incorporating a few rather accessible measures into routine clinical monitoring. Features whose recording can result in better predictability are shown in bold in Figure 3. Indeed, since the challenge ran, clinics involved in the Italian registry used in this challenge have been careful to add measurements of patient weight, BMI and respiratory functions.

#### Clinically relevant patient clusters

In this challenge we used a crowdsourcing approach to explore different stratification schemes for ALS patients and use them to identify clinically significant patient sub-populations. Such sub-populations can be of great value for understanding the ALS pathology, improving clinical care and planning better, more efficient clinical trials. We were also hoping to learn whether clustering could help improve the performance of prediction algorithms, especially when applied to heterogeneous patient populations (i.e., registry data) when the number of available clinical features is limited.

Most challenge participants chose not to incorporate a clustering component in their prediction algorithm, and overall, we did not observe any consistent advantage (in terms of improved prediction accuracy) for clustering in any of the evaluation metrics. The main goal of clustering, however, was not necessarily to impact prediction accuracy, but to reveal, across a large variety of methods, consistent clusters of patients that could uncover clinically significant ALS patient subpopulations. This sort of analysis can only be accomplished in the context of a large communal effort that allows comparison of independently generated algorithms working on sufficiently large datasets, as was the case in this challenge. To identify consistent clusters, we assigned patient pairs with scores reflecting how often they appear together in the same cluster across all different algorithms and used the resulting connectivity matrix as input to correlation-based k-means analysis (see online methods and Supplementary material part 6). The resulting clusters reflected groups of patients that consistently and significantly tend to be clustered together across different clustering schemes and algorithms. The different clustering methods led to substantially different results, for instance, in the registry progression and survival subchallenges the numbers of clusters varied from 2 to 131 and from 2 to 159, respectively.

An important result of this analysis was that the patients of each of the four sub-challenges could be easily clustered into in a consistent small set (3-4) of visually discrete consensus clusters (Figure 4). In both sub-challenges based on PRO-ACT, patients within consensus clusters were significantly more strongly connected than expected by chance (pairs of patients co-clustered significantly with FDR<5% are depicted as edges in Figure 4b). The stronger clustering effect in the PRO-ACT vs. the registry sub-challenges is likely a reflection of sample size (10,000+ vs. ~1,500 patients, respectively).

**Figure 4:**
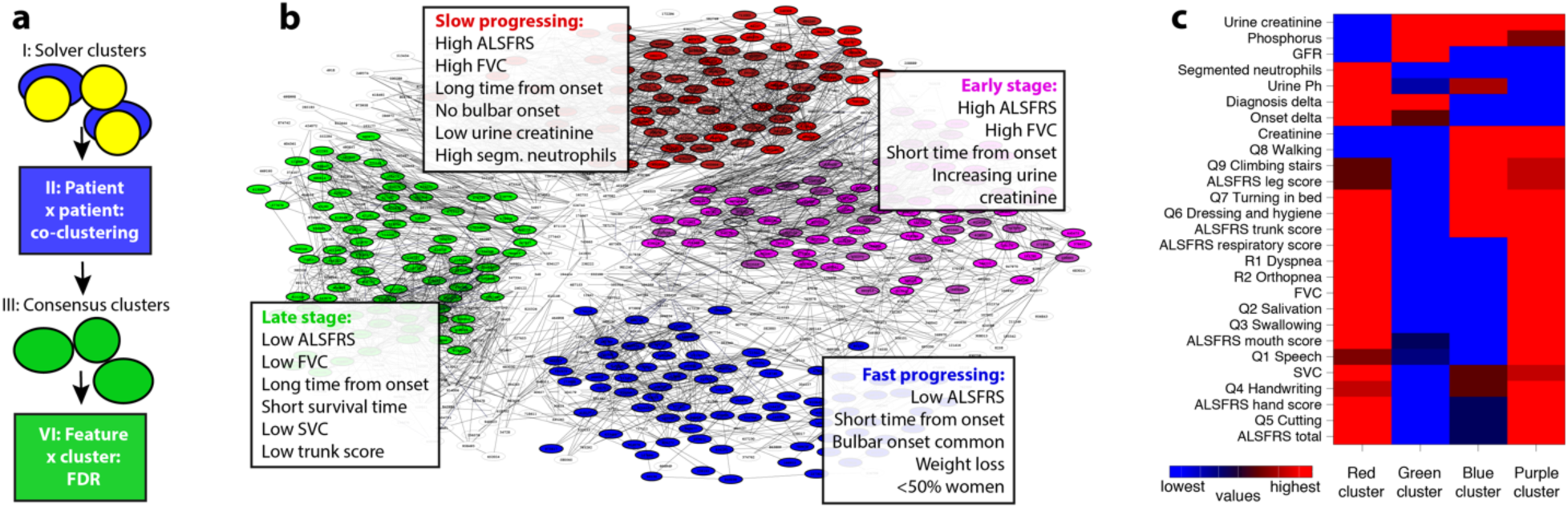
Overview of the consensus clustering. (a) Outline of consensus clustering method: the tendency of patients to co-cluster was assessed across cluster-sets generated independently by the different solvers (I). The resulting connectivity matrix (II) was then used as input for obtaining consensus clusters by k-means (III). Finally, False discovery rates (FDRs) were estimated by ANOVA to assess, which features were differentially distributed between consensus clusters (IV). (b) Graph-based clustering of the connectivity matrix for the PRO-ACT progression sub-challenge. Nodes in the graph represent patients that are colored based on their k-means cluster, if they correspond to the 50% of patients closest to their respective cluster centroid. Edges denote pairs of patients with a significant chance of being co-clustered. (c) We compared features (names starting with Q/R are ALSFRS component scores from the original or revised scale, respectively) between pairs of clusters (columns in heatmap) by t-tests/FDRs. Different colors within heatmap rows indicate values that are significantly different between clusters (FDR < 5%) on the scale from the lowest (blue) to highest (red). Notable results are listed in Panel b.

Overall, consensus clusters could be broadly regarded as classifying patients as slow progressing (“red”), fast progressing (“blue”), early stage (“purple”) or late stage (“green”). We chose to focus on clinical characteristics of the PRO-ACT progression sub-challenge, since these consensus clusters reached the highest level of statistical significance, but similar clusters were found across all sub-challenges.

One cluster, the “red” or “slow progressing” cluster included patients who, despite having experienced symptoms for a relatively long time (2.2 years from disease onset, on average) still maintained relatively high functional capabilities (average ALSFRS-R scores of 40.25 at the beginning of the clinical observation period), with functional impairments mostly limited to limbs (either legs or arms) and little bulbar or respiratory involvement. Accordingly, these patients had slow disease progression (annual average loss of −0.48 on the ALSFRS-R scale). These are the patients with the best prognosis for ALS and therefore noteworthy to assess closely as they might aid clinicians in predicting favorable disease progression. First, only few (3%) of these “slow progressing” patients had bulbar onset. While bulbar onset had been frequently correlated with poorer prognosis^22-24^, this cluster analysis suggests that bulbar patients will rarely be classified as a slow progressing disease presentation. A second important observation is low creatinine levels (average of 67 mcmol/L in the first three month of data collection, compared to a normal desired range of 74.3-107mcmol/L). Creatinine was reported in the previous challenge^16^, as a predictor of disease progression and these results suggest that it might also serve as a predictor of this special case of patients with improved prognosis.

A very similar cluster was observed in the analysis of clusters derived from the PRO-ACT survival sub-challenge, with patients living with ALS for an average of 1.5 years while displaying little functional decline (average total ALSFRS_R score 39.4), largely intact breathing functionality (FVC 94.4% of normal on average) and an ALSFRS progression rate of −0.72points/months. To characterize clusters and the involved patients for clinical relevance, we compared all pairs of clusters (using ANOVA and t-test, resulting in multiple-testing corrected false discovery rates or FDRs) to assess which features had values specifically different between the clusters (Figure 4c). We also examined the correlation between feature values and clinical outcome (progression rate) in each cluster to identify features which were important for prediction in some clusters but not in others. Based on the clusters clinical features analysis, the two features unique for this “slow progressing” cluster were urine creatinine and segmented neutrophils (Figure 4b-c). Both of these features are therefore potential biomarkers of slow progression in ALS. Neutrophils were indeed found to be connected to ALS progression in some studies^33,34^ but not others^35^ and the stratification to sub-groups might shed further light on these results.

Another cluster that was superficially similar but in fact quite different was the “purple” or “early stage patients”. In our pair-wise comparison of features across clusters, the “early stage” and “slow progression” clusters (“purple” and “red” clusters respectively) were similar in having high ALSFRS and FVC scores in the first 3 months of assessment, indicating little functional impairment (Figure 4b-c). However, the distinctive feature of patients in the “purple” cluster was the fact that they were early in their disease, on average ~10 months from symptom onset. Thus, the largely preserved functional state of this patients could be attributed to their early disease stage, rather than a slow progression rate. Indeed, with time these patients became fast progressors (−0.93 ALSAFRS-R points decline monthly on average) and had marginally lower than normal creatinine levels (72.25 mcmol/L on average on the first three months of data). Curiously, urine creatinine was correlated with disease progression (r^2^=0.467 p=0.01, see Supplementary material 7 for figures) only for this cluster, suggesting again that urine creatinine might be useful predictor of disease progression already early in the disease. The relationship between Urine creatinine and serum creatinine and with both to muscle breakdown is not straight forward^36^.

A third “green” cluster of “late stage” patients included patients that were clinically late in their disease (although not late chronologically- on average 1.7 years from disease onset), who were severely disabled in all functional domains (average ALSFRS-R score of 29 points at the beginning of the clinical observation period) and were displaying early signs of breathing dysfunction (average FVC 77%) and shorter survival time (1.5 years on average).

These patients had a significant correlation between their ALSFRS “trunk” score (questions about dressing and hygiene and turning in bed) and disease progression (r^2^=0.314, p=0.001), which might be indicative of their advanced disability status. The comparison of feature values across clusters revealed two features which were uniquely predictive only for this cluster: Slow Vital Capacity (SVC), and measurements of “trunk” ALSFRS scores (Figure 4c, examples are available in Supplementary material part 7). These two classifiers should be taken into consideration clinically as they might be stronger indicators that the patients are reaching the final stages of their disease. The “trunk” score indeed represents complex functions that require the combined efforts of upper and lower motor neurons, and is therefore more clearly impaired later in the disease. SVC, while highly correlated with FVC, might became predictive later in the disease when respiratory function diminishes. They could also be used for clinical trial exclusion criteria to improve patient survival throughout the trial. Indeed, SVC was recently suggested as an indicator of respiratory failure in ALS^37^.

The last cluster, “blue” or “fast progressing” patients, represents the most critical patients, who have been experiencing symptoms for only 10 months on average, but were already significantly impaired in all functional categories (average ALSFRS-R score of 35.75) at the beginning of the clinical observation period. These patients continue to have a very fast disease progression rate (ALSFRS progression slope of –1.05points/month) and on average die 662 days after the start of clinical observation, amounting to an overall average survival of only 2.7 years from disease onset. This is the population of the most urgent need and merits a closer investigation. Half of the patients in this cluster had bulbar onset (compared to 20% bulbar onset across all patients) and were more likely to have lower scores in all ALSFRS_R functional domains (leg, hand, trunk, bulbar and respiratory functions) and to have significant weight loss over the 1-year observation period (average of 5kg lost per year).

Importantly, a similar yet more severe cluster was observed in the PRO-ACT survival data, with patients showing diminished disease states (initial ALSFRS-R of 24 ALSFRS points) and survival (average of 443 days from trial onset). When clustering for survival, it was noticeable that women are more likely to be fast progressors (53% women compared to 40% in the general patient population), even beyond their higher likelihood to have bulbar onset (32% of women). This cluster of patients with the poorest prognosis was also found in the registry consensus clustering for both the progression and the survival data, with short survival times, fast progression and high rate of bulbar onset patients.

There were also features that stood in our pair-wise comparison because they were significantly different between almost all pairs of clusters. Unsurprisingly the most discriminative feature was time of onset (which was significantly different between the “red”, “blue” and “green” clusters). However, equally discriminative were also ALSFRS question 1 (speech) and our combined measure mouth, averaging ALSFRS questions 1-3 (speech, swallowing and salivation), highlighting again the important role of bulbar function in discriminating ALS consensus clusters.

## Discussion

Disease heterogeneity, and particularly large unexplained variance in disease progression rate and survival, is a hallmark feature of ALS disease^38^. The range of ALS clinical course includes patients such as Lou Gehrig, who perished within 2-3 years of symptoms onset, to patients such as Stephen Hawking, who has been living with the disease for several decades. The hypothesized existence of distinct subgroups of ALS patients and their importance for ALS research and clinical care were highlighted in recent years by clinical trials in which only a subset of patients responded to the tested treatment^3,39^. Despite some recent advances from genetic studies^40,41^, there is currently no generally accepted stratification scheme for ALS patients, and more importantly, there no way of using stratification information for tailoring survival and disease progression estimates specifically to individual patients. The ALS Stratification Challenge was a global crowdsourcing effort aimed to develop new tools and insights for understanding patient subpopulations as they relate to ALS disease progression and survival. Thirty teams from around the world submitted algorithms to the challenge, with the winning solutions outperforming currently available prediction algorithms. The ALS Stratification Challenge followed the success of the previous ALS Prediction challenge^16^, and adapted versions of the best performing algorithms from the previous challenge were used as baseline methods against which the new algorithms were evaluated. The current challenge included additional data and design features that made its resulting algorithms more robust and more relevant for clinical application, including the use of community-based data, the limitation on the number of features used for prediction, and the prediction of survival as well as disease progression. The current challenge added another requirement highly relevant to the application for future clinical trials: the validation of algorithms on a dataset derived from completely independent clinical trials. We used the prediction and stratification algorithms submitted to the challenge to suggest a new stratification scheme for ALS patients which includes detailed characterization of patient sub-populations as well as highlighting the most relevant clinical features for classification and outcome prediction for each patient group.

The current challenge invited participants to develop prediction algorithms either based on clinical trial data (from the PRO-ACT database) or from ALS registries containing data collected through ALS clinics. While PRO-ACT data might be particularly useful for developing prediction algorithms to be used in clinical trials, registry data should be more clinically relevant and include a more accurate and complete representation of the entire patient population^42^. This is the first time that registry data was made publicly available, and the design of the challenge enabled us to directly compare performance of prediction algorithms when applied to PRO-ACT vs. registry data. The smaller volume of available data in the registries dataset, both in terms of number of patients and of number of clinical features recorded per patient, combined with the increased heterogeneity of patients^43^, made prediction more challenging and overall registry prediction algorithms achieved lower levels of prediction accuracy. We suggest a number of clinical features (Fig. 3), such as FVC and weight, which could be added to routine clinical assessment to potentially improve prediction accuracy and aid clinicians in predicting individual patient prognosis. Conversely, a number of features that could be found exclusively in the registries data, including common genetic mutation data and detailed onset site assessments, were both highly informative for prediction and should be considered for incorporation into ALS clinical trial screening or baseline assessment.

We examined the features most often chosen for prediction by the different challenge participants to see what features had the highest predictive power. This analysis revealed several features, already well reported in the literature, such as age, gender^22–24^ and respiratory capacity.^22,29,44,45,46^ which were strong predictors of survival, but were less informative for predicting disease progression, whereas other features, including limb motor function ALSFRS-R scores (hands and legs function) and specific ALS staging scores^47^ were more informative for predicting disease progression rather than survival. Creatinine, already suggested before as a measure predictive of ALS prognosis^16,29,48,49^,^50^ was also found to be predictive in this challenge, and interestingly this was specific for patients early in their disease. Segmented neutrophils were also suggested by our analysis as a relevant novel predictor, specifically for slower progressing patients, while SVC and ALSFRS “trunk” scores were associated with outcomes only specifically for patients later in their disease.

The main goal of this challenge was to uncover clinically meaningful subgroups of ALS patients, a challenging task since no known “gold standard” exists for ALS patient stratification. In this study we applied a “bottom-up” approach to patient stratification: we did not make any a-priory assumptions regarding patient sub-populations, but instead defined patient clusters by a “consensus vote” based on participants’ submitted algorithms. Challenge participants were free to base their clustering on any subset of the available clinical data, choose any type of clustering method and any number of clusters. While clustering was not directly related to any immediate benefit in algorithms’ prediction accuracy, it did reveal consistent patterns of patient classification that are of great potential interest. We suggest that these clusters could be used to identify subgroups of patients to guide further research of disease mechanisms and the planning of individual patient care programs and ALS clinical trials. Given the covert nature of ALS patient stratification, only a large-scale crowdsourcing effort, where different and independent teams apply diverse methods on a similar and large enough dataset can uncover such an underlying population structure free from a priory assumptions.

This communal approach indeed revealed a few sub-groups of patients which not only tended to cluster together across different algorithms but also displayed similar characteristics across different sub-challenges - clusters which may be the basis for a new stratification framework for ALS patients. Overall, we observed four patient groups: slow progressing patient and fast progressing patients, as well as patients with an average progression rate which were either early or late in their disease at the beginning of the recorded clinical observation period. The wealth of lab tests available in the PRO-ACT dataset also uncovered a few clinical tests characteristic to each of these subgroups (urine creatinine and segmented neutrophils), but it is currently unclear to what extent these could be used as predictive markers for patient stratification or are merely a reflection of the patients’ medical condition. However, our results do suggest that a few easy tools, such as early testing for FVC and creatinine could improve prediction of individual patient prognosis, for example, helping to differentiate slow and fast progressing patients as early as possible in their diseases.

Overall, these results suggest a novel stratification scheme for ALS, with relevant classifiers and group-specific predictors, and it would be highly interesting to further prospectively assess its benefit for clinical care and clinical trials. As most clinical trials enroll patients with high functions, they might over-represent patients from the “early” patient” and “slow progressing” cluster and under-represent the other two cluster of patients. It would also be interesting to assess whether the predictors revealed here can uncover new insights about disease mechanisms.

To summarize, the ALS Stratification Challenge and subsequent analysis provided us with algorithmic tools and insights about stratifying ALS patients and uncovered meaningful sub-groups of patients. Patient stratification is always a challenge, and even more so for rare diseases with substantial heterogeneity where effective stratification can advance clinical development substantially. Stratification is also highly needed for clinical care, to allow more personalized treatment. The tools and insights presented in this study can offer a first attempt for improvement in clinical trial development, selection and interpretation, accelerating the development of a much-needed treatment for a dire aliment such as ALS.

## Supplementary information

We provide additional method description and results (ALS Stratification Supplement.pdf), the solver-provided descriptions of submitted algorithms (Challenge writeups.pdf) as well as data on solver predictions, the cluster analysis and assessment scripts (stratification.zip, see readme files contained there for details).

### Acknowledgements

We are grateful to the following people for their important assistance with this manuscript: The clinicians and researchers behind the Irish and Italian ALS registers and the pharmaceutical companies which provided data to the PRO-ACT dataset, that enabled this entire endeavor, the hundreds of participants on the crowdfunding effort that provided this challenge’s award. Prof. David Schoenfeld for his assistance with statistical considerations, and, of course, the solvers who participated in the challenge and the patients who inspired this effort.

Data used in the preparation of this article were obtained from the Pooled Resource Open-Access ALS Clinical Trials (PRO-ACT) Database. As such, the following organizations and individuals within the PRO-ACT Consortium contributed to the design and implementation of the PRO-ACT Database and/or provided data, but did not participate in the analysis of the data or the writing of this report: Neurological Clinical Research Institute at MGH, Northeast ALS Consortium, Novartis, Prize4Life Israel, Regeneron Pharmaceuticals Inc., Sanofi, Teva Pharmaceutical Industries Ltd.

## AUTHOR CONTRIBUTIONS

R.K., N.Z., R.N., T.N, L.M, G.S. and M.L.L. designed the challenge. O.H, M.C and A.C contributed and prepared the data and providing clinical insight, R.K., B.D, J.G-G, R.N, L.W and G.L prepared the data and baseline algorithm, J.G-G, R.K, B.H, V.B, J.K and D.D provided challenge data and technical support, N.Z. and R.K. managed the challenge execution. R.K., M.B, R.N and N.Z. analyzed the results and wrote the paper. The ALS Stratification Consortium provided methodology. S.J. edited and reviewed portions of the paper.

## Online Methods

### Datasets

The challenge made use of two datasets: data collected during clinical trials (PRO-ACT) and data collected during routine visits to ALS clinics (ALS registries). Both datasets included both static and time resolved measurements covering a wide range of data types and clinical measurements (full data dictionaries for both datasets can be found in the supplementary material). The time in which measurements were taken was noted in days relative to clinical trial onset or to first clinic visit available on records (time “delta”). Data was provided to challenge participants in tabular form. Each line represented the measurement of a single feature for a single patient at a particular time point Outcome measures (ALSFRS slope for progression and survival) for the training datasets were provided to challenge participants in separate tables.

#### PRO-ACT

The Pooled Resource Open-Access ALS Clinical Trials (PRO-ACT) Database was formed in 2011 by Prize4Life in collaboration with the Neurological Clinical Research Institute (NCRI) of Massachusetts General Hospital, the Northeast ALS Consortium, and with funding from the ALS Therapy Alliance. PRO-ACT contains data collected during phase II/III ALS clinical trial, volunteered by PRO-ACT Consortium members. Data from 17 clinical trials (Atassi et al 2014) was used for the previous prediction challenge [Kueffner et al, 2015] and later made publicly available via the PRO-ACT web platform (www.ALSdatabase.org). This data was used for algorithms’ training in the current challenge. Data from 6 additional clinical trials (Cudkowicz et al 2013, Cudkowicz et al 2014, Gordon et al 2007, Kaufmann et al 2009, Sorenson et al 2008, Swati et al 2010), never before made publicly available, was homogenized to comply with PRO-ACT data format standards and used for algorithms’ validation and assessment. Similar to the design of the previous challenge, the data used for validation was randomly subdivided into a test set (leaderboard data, 400 patients) and a validation set (1,488 patients). Challenge participant could submit versions of their code to be tested on the test set (limited to once per week per team to avoid overfitting) with the results published on a public leaderboard which served to provide feedback to participants. After the challenge was completed all submitted algorithms were assessed using the validation set data (Table 1).

#### ALS registries data

Community-based data, never before publicly released, used in this challenge was collected through two ALS registries: 1) The Irish National ALS Register including data collected from ALS clinics in Ireland. 2) The Piemonte and Valle d’Aosta Register for ALS, including data collected from ALS clinics in Piemonte and Valle d’Aosta region of Italy. Data from the two registries was merged, harmonized and converted to the same format as the PRO-ACT data. Data was divided into a training (986 patients) and a validation (493) set. Stratification of data ensured a similarly proportional representation of patients from the Irish and Italian cohorts but was otherwise done randomly. Due to the small number of patients available in this dataset there was no test set (leaderboard data) available to challenge participants.

### Definition of predicted outcome measures - “gold standard” calculation

#### Disease progression

ALS disease progression was defined as the slope of the total ALSFRS score, similar to the definitions used in the 2012 prediction challenge [Kueffner et al. 2015]. Briefly, ALSFRS was calculated as:

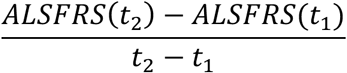

Where t_1_ and t_2_ were the time points at the first and last clinic records in the relevant time period 92-365 days in which total ALSFRS scores were recorded. Whereas time data in the both challenge datasets was given in days, it was converted to months for the calculation of ALSFRS slope, according to the following: t_(months)_= (t_(days)_/365) *12). Patients had to have at least two clinical records in the relevant time period for their data to be used for validation (see Table 2). Participants were required to write algorithms that would predict ALSFRS slope between 3 and 12 months, based on data collected in the first three months of clinical records.

#### Survival

Survival was defined as time until death or until tracheostomy surgery (the introduction of invasive breathing tube-time where without intervention the patient was unlikely to survive), whenever this information was available. For patients who had no time of death logged on the clinical records the time of the last clinical visit was recorded in the survival records with a status indicating they were alive.

Patients whose final records were on or before day 90 (from onset of clinical trial or of clinical records) were excluded from the survival analysis (see Table 2). Challenge participants were required to write algorithms that predicted the likelihood of survival for each patient at three time points; 12, 18 and 24 months from the onset of clinical records.

#### Predictions assessment and scoring scheme

All methods for performance assessment were based on evaluating how close the algorithms’ predictions were compared to the gold standards, averaged across all patients. We used three different evaluation metrics to assess submitted algorithms’ performance in the disease progression sub-challenges. Two methods, the root mean square deviation (RMSD) and Pearson’s correlation (PCC) were used and described in the previous ALS prediction challenge (Kueffner et al., 2015).

In the current challenge we added a third evaluation metric: the concordance index (CI), which evaluates the similarity of ranks between predicted and actual ordered lists of measurements. The concordance index was the only metric used to evaluate performance in the survival prediction sub-challenge, since it is commonly used in survival analysis and is best suited for assessing censored data [Noren et al., 2016]. When there is no censored data or ties, the *c*-index between a predicted list of survival times of *n* patients, *Pred* = {*p_1_, p_2_*,…, *p_n_},* and the actual survival times for the same *n* patients, *Actual* = {*a_1_,a_2_,…, a_n_*}, is calculated as:

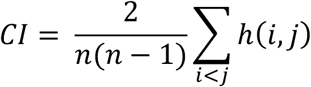

where,

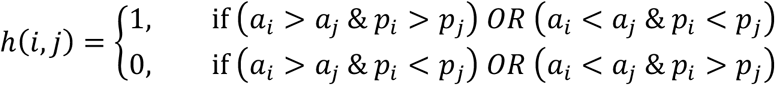

Please see supplementary data on details accommodating for the possibilities of ties and censored data.

The three performance assessment (or three CI values, in the case of survival prediction) were combined using Z-score transformation. We generated 100,000 random sets of predictions for each of the four sub-challenges, by randomply assigning “gold standard” progression or survival values to patients. We then calculated the RMSD, PCC and CI scores for each of these random shuffles, and used the resulting three sets of 100,000 values to calculate the mean and standard deviation (SD) of each of the three scoring metrics. These mean and SD values were used to calculate the Z-scores for the three assessment metrics for each of the submitted algorithms, given by:

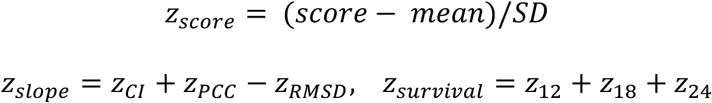

Outcomes of the survival sub-challenges were further validated by a receiver operating characteristic analysis (time ROC analysis). For the purpose of the challenge, time ROC was calculated using the R-package timeROC (https://cran.r-project.org/web/packages/timeROC/timeROC.pdf). On the plus side, the timeROC analysis incorporated the three survival time frames to be predicted (12, 18, 24 months), and performance is thus specific to the time frame in contrast to the analysis via CI, where the same set of predictions would evaluate the same irrespective to the time frame. However, the CI is better suited to represent the right censured nature of the survival data. The rank ordering of submitted algorithms for both survival sub-challenges was very similar when comparing the outcomes of the combined z-transformed CI index and the time ROC analysis. Results of the time ROC analysis could be found in the supplementary material.

### Consensus clusters and determination of discriminating features

For each subchallenge, we integrated the teams’ clusterings into a consensus clustering. First, we created a square patient x patient co-clustering matrix *M* where each matrix element *m*_*i,j*_ contained a normalized score that expressed how often the corresponding patients *i* and *j* appear together in a single cluster across all team submissions. The normalized score takes the size of the teams’ clusters into account such that two patients clustered together in a smaller cluster receive a larger weight. The scores *m*_*i,j*_ were calculated as the sum of contributions (eq. 1), across all submissions *s*, where patients *i* and *j* appear together in the same cluster:

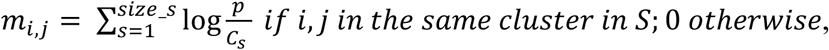

where *p* is the total number of patients in the given sub-challenge and *c*_*s*_ is the number of patients in the cluster of submission *s* that includes both *i* and *j* (to normalize by cluster size).

We next determined the statistical significance of *m*_*i,j*_ values, i.e. to determine whether patients *i* and *j* are clustered together more often than expected by chance. We randomly assigned patients into clusters of the same size as contained in the original submissions, and repeated this random sampling process 100 times. We then calculated *m*_*i,j*_ scores for each set of randomly assigned clusters, giving us 100 *m*_*i,j*_ scores for each *i,j* patient pair. We used these permuted *m*_*i,j*_ value to calculate a false discovery rate for any given *m*_*i,j*_ score derived from the participants’ submissions data, by assigning:

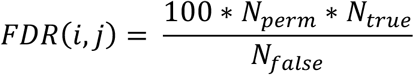

where *N*_*perm*_ is the number of permutations, *N*_*true*_ and *N*_*false*_ refer to the number of scores computed from submitted and permuted data, respectively, that were less or equal than the given score. This approach to FDR calculation was adapted from significance analysis of microarrays (Tusher et al., 2001) and described in detail in previous work (Marozava et al., 2014). Subsequently a FDR threshold of 5% was applied to identify significantly pairs of patients that significantly tend to be clustered together.

A graph of significant pairs was then plotted using the graphviz software package in order to visually determine a plausible number of clusters k. To generate the final patient clusters, we performed k-means clustering for *k* clusters based on the matrix M using average linkage and Pearson’s correlation metric. Thus, the correlation metric calculates the distance between patients *i* and *j* by comparing the corresponding rows, denoted as *m_i_.* and *m_j_.,* in the matrix *M.* This analysis identified three consensus clusters for both survival and the registry progression sub-challenges and four clusters for the PROACT progression sub-challenge.

### Detection of features differentially distributed across patient groups

After finding a small number of consensus clusters of patients for each sub-challenge, we determined the features that discriminate between consensus clusters by statistical tests. Values of longitudinal variables (e.g., ALSFRS) were averaged across time, and we looked at average values of two different time periods consistent with the challenge’s overall design: 0-3 months and 3-12 months. We applied one-way analysis of variance (ANOVA) to continuous features (e.g. age of onset) and Fisher’s exact test to discrete features (e.g. site of onset). This leads to two matrices, *C* and D, capturing the values of the continuous and discrete features, respectively. Rows in these matrices correspond to the features, columns to the patients. In addition, each feature vector (containing the values for that feature across the *p* patients) was randomly permuted a hundred times and statistical tests were applied to the 100 randomized datasets to compute FDRs (analogous to the description above) separately for the results from Fisher’s exact test and ANOVA.

We then regard the relative ranks from the ANOVA and Fisher as new test values. From these, we compute FDRs now across discrete and continuous features together. By this integrated statistical assessment we obtained FDR values for all features. Features were deemed statistically significant across all clusters if they exhibited FDR values of <5%. See below pseudo code for details.

Subsequently, t-tests were applied to identify all pairs of clusters where ANOVA determined that continuous features exhibited significantly differential values (FDR < 5%). The same permutations were applied to the feature vectors as above, such that we were able to transform t-test p-values into false discovery rates analogously. As before, we regard a feature as statistically significantly different between two clusters if such a comparison resulted in an FDR <5%. FDRs do not need multiple testing correction as it is already built into the permutation test. Note that we applied the ANOVA “trick” here: pairwise comparisons were only performed (and thus subject to multiple testing correction) for features with overall significance across all clusters. This leads to a less severe multiple testing correction and accordingly to a more sensitive test for the pairwise comparisons in contrast to the case were pairwise comparisons would have been performed and corrected for all features.

We then transformed the pairwise comparisons for each feature into cluster specific levels *l* in order to create a heatmap. If a comparison between a pair of clusters *(a,b)* is significant, and the given feature displays higher values in *a* as compared to *b,* we applied *l(a)=l(a)+1* and *l(b)=l(b)−* 1. After all pairwise comparisons are integrated this way, the scale of all features was linearly transformed into a range between [−1 … +1]. Thereby, the heatmap clearly displays for each feature the distinguishable cluster levels.

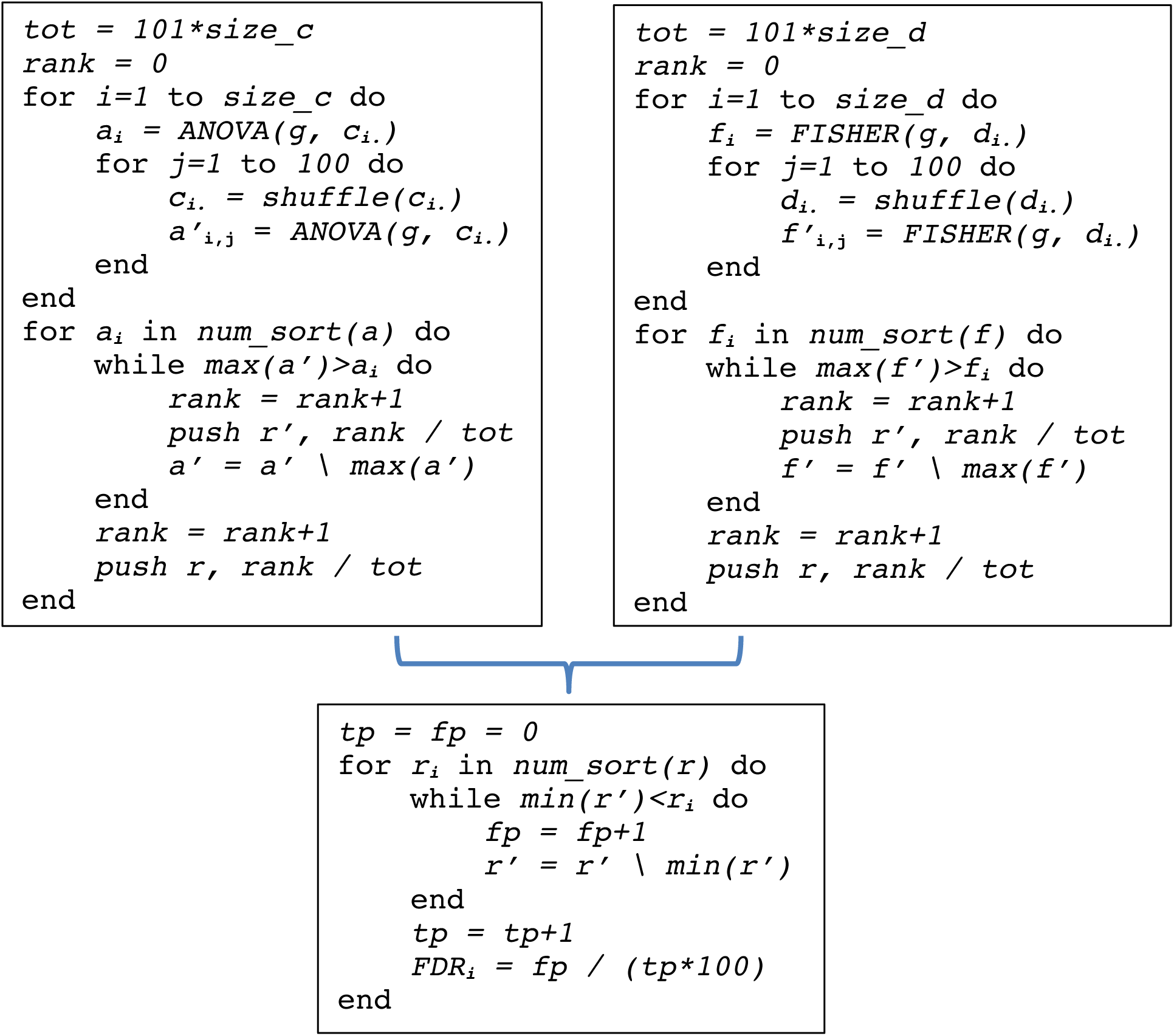

### Pseudo code analysis of differentially distributed features

The pseudocode in the left panel illustrates the computation of the ANOVA test statistic *a* at the example of the continuous features in C. The design *g* specifies the mapping of patients to clusters. A permuted test is calculated by shuffling values in rows of the matrix C 100 times, computing their associated test statistics *a’* and pushing their relative ranks and the relative ranks of *a* separately into arrays *r’* and *r,* respectively. The backslash notation denotes the removal of an element, i.e. in a’=a’ \ max(a’), the entry with the highest value is removed from a’. Analogously, the FISHER test statistic *f* is calculated for the discrete features in *D* (pseudo code in the right panel). Finally, FDRs are calculated by comparing the relative ranks from the true statistics r vs. the relative ranks from the permuted statistics *r’* across discrete and continuous features (lower panel).

## References

1 Swinnen, B., Robberecht, W. The phenotypic variability of amyotrophic lateral sclerosis. Nat Rev Neurol. 10,661–70 (2014).

2 Miller, R.G., Mitchell, J.D., Lyon, M., Moore, D.H., Riluzole for amyotrophic lateralsclerosis (ALS)/motor neuron disease (MND). Cochrane Database Syst Rev. CD001447 (2002).

3 Edaravone (MCI-186) ALS Study Group. Safety and efficacy of edaravone in well defined patients with amyotrophic lateral sclerosis: a randomised, double-blind, placebo-controlled trial. Lancet Neurol. 16, 505–512 (2017).

4 Ravits, J.M., La Spada, A.R. ALS motor phenotype heterogeneity, focality, and spread: deconstructing motor neuron degeneration. Neurology. 73, 805–11 (2009).

5 Logroscino, G. Classifying change and heterogeneity in amyotrophic lateral sclerosis. Lancet Neurol. 15, 1111–2 (2016).

6 Kenna, K.P., et al. Delineating the genetic heterogeneity of ALS using targeted high-throughput sequencing. J Med Genet. 50, 776–83 (2013).

7 Turner, M.R. et al. Controversies and priorities in amyotrophic lateral sclerosis. Lancet Neurol. 12, 310–22 (2013).

8 Sabatelli, M., Conte, A., Zollino, M. Clinical and genetic heterogeneity of amyotrophic lateral sclerosis. Clin Genet. 83, 408–16 (2013).

9 Brooks, B.R. El Escorial World Federation of Neurology criteria for the diagnosis of amyotrophic lateral sclerosis. Subcommittee on Motor Neuron Diseases/Amyotrophic Lateral Sclerosis of the World Federation of Neurology Research Group on Neuromuscular Diseases and the El Escorial “Clinical limits of amyotrophic lateral sclerosis” workshop contributors. J Neurol Sci. 124, Suppl:96–107 (1994).

10 Carvalho, M.D., Swash, M. Awaji diagnostic algorithm increases sensitivity of El Escorial criteria for ALS diagnosis. Amyotroph Lateral Scler. 10, 53–7 (2009).

11 Ganesalingam, J., et al Latent cluster analysis of ALS phenotypes identifies prognostically differing groups. PLoS One. 4, e7107 (2009).

12 Su, X.W., et al. Biomarker-based predictive models for prognosis in amyotrophic lateral sclerosis. JAMA Neurol. 70, 1505–11 (2013).

13 Elamin, M., Predicting prognosis in amyotrophic lateral sclerosis: a simple algorithm. J Neurol. 262, 1447–54 (2015).

14 Marin, B., et al. Stratification of ALS patients’ survival: a population-based study. J Neurol. 263, 100–11 (2016).

15 Atassi, N., et al. The PRO-ACT database: design, initial analyses, and predictive features. Neurology. 83, 1719–25 (2014).

16 Küffner, R., et al. Crowdsourced analysis of clinical trial data to predict amyotrophic lateral sclerosis progression. Nat Biotechnol. 33, 51–7 (2015).

17 Zach, N., et al. Being PRO-ACTive: What can a Clinical Trial Database Reveal About ALS? Neurotherapeutics. 12, 417–23 (2015).

18 Taylor, A.A., et al Pooled Resource Open-Access ALS Clinical Trials Consortium. Predicting disease progression in amyotrophic lateral sclerosis. Ann Clin Transl Neurol. 3, 866–875 (2016).

19 Norel, R., Rice, J.J., Stolovitzky, G. The self-assessment trap: can we all be better than average? Mol Syst Biol. 7, 537 (2011).

20 Marbach, D., et al. Wisdom of Crowds for Robust Gene Network Inference. Nature methods 9, 796–804 (2012).

21 Rhrissorrakrai, K., et al. Understanding the limits of animal models as predictors of human biology: lessons learned from the sbv IMPROVER Species Translation Challenge. Bioinformatics. 31, 471–83 (2015).

22 Huang, Z., et al. Complete hazard ranking to analyze right-censored data: An ALS survival study. PLoS Comput Biol 13, e1005887 (2017).

23 Magnus, T. et al. Disease progression in amyotrophic lateral sclerosis: predictors of survival. Muscle Nerve 25, 709–714 (2002).

24 del Aguila, M., Longstreth, W., McGuire, V., Koepsell, T. & Van Belle, G. Prognosis in amyotrophic lateral sclerosis: a population-based study. Neurology 60, 813–819 (2003).

25 Pastula, D.M. et al. Factors associated with survival in the national registry of veterans with ALS. Amyotroph. Lateral Scler. 10, 332–338 (2009).

26 Czaplinski, A., Yen, A.A., Appel, S.H. Amyotrophic lateral sclerosis: early predictors of prolonged survival. J Neurol. 253, 1428–36 (2006).

27 Paganoni, S. et al. Uric acid levels predict survival in men with amyotrophic lateral sclerosis. J. Neurol. 259, 1923–1928 (2012).

28 Chiò, A., et al. Piemonte and Valle d’Aosta Register for Amyotrophic Lateral Sclerosis. Amyotrophic lateral sclerosis outcome measures and the role of albumin and creatinine: a population-based study. JAMA Neurol. 71, 1134–42 (2014).

29 Pinto, S., Gromicho, M., de Carvalho, M. Sialorrhoea and reversals in ALS functional rating scale. J Neurol Neurosurg Psychiatry. 88, 187–188 (2017).

30 Paganoni, S., Deng, J., Jaffa, M., Cudkowicz, M.E. & Wills, A.M. Body mass index, not dyslipidemia, is an independent predictor of survival in amyotrophic lateral sclerosis. Muscle Nerve 44, 20–24 (2011).

31 Paganoni, S., Deng, J., Jaffa, M., Cudkowicz, M.E. & Wills, A.M. What does body mass index measure in amyotrophic lateral sclerosis and why should we care? Muscle Nerve 45, 612 (2012).

32 Corcia, P., et al. Causes of death in a post-mortem series of ALS patients. Amyotroph Lateral Scler. 9, 59–62 (2008).

33 Murdock, B.J. et al. Increased ratio of circulating neutrophils to monocytes in amyotrophic lateral sclerosis. Neurol Neuroimmunol Neuroinflamm. 3, e242 (2016).

34 Murdock, B.J., et al. Correlation of peripheral immunity with rapid amyotrophic lateral sclerosis progression. JAMA Neurol. (2017).

35 Chiò, A et al. Amyotrophic lateral sclerosis outcome measures and the role of albumin and creatinine: a population-based study. JAMA Neurol. 71, 1134–42 (2014).

36 Baxmann, A.C. et al. Influence of Muscle Mass and Physical Activity on Serum and Urinary Creatinine and Serum Cystatin C. Clinical Journal of the American Society of Nephrology?: CJASN. 3:348–354 (2008).

37 Andrews, J.A et al. Association between decline in slow vital capacity and respiratory insufficiency, use of assisted ventilation, tracheostomy, or death in Patients with amyotrophic lateral sclerosis. JAMA Neurol. (2017).

38 Bedlack, R.S., et al. How common are ALS plateaus and reversals? Neurology. 86, 808–12 (2016).

39 Fiala, M., Mizwicki, M.T., Weitzman, R., Magpantay, L., Nishimoto, N. Tocilizumab infusion therapy normalizes inflammation in sporadic ALS patients. Am. J. Neuro. Dis. 2,129–139 (2013).

40 Fogh. I., et al. Association of a Locus in the CAMTA1 Gene with Survival in Patients With Sporadic Amyotrophic Lateral Sclerosis. JAMA Neurol. 73, 812–20 (2016).

41 Umoh, M.E., et al. Comparative analysis of C9orf72 and sporadic disease in an ALS clinic population. Neurology. 87, 1024–30 (2016).

42 Chiò, A. et al. ALS clinical trials: do enrolled patients accurately represent the ALS population? Neurology 77, 1432–1437 (2011).

43 Rooney, J.P.K., et al. Benefits, pitfalls, and future design of population-based registers in neurodegenerative disease. Neurology. 88, 2321–2329 (2017).

44 Traynor, B., Zhang, H., Shefner, J., Schoenfeld, D. & Cudkowicz, M. Functional outcome measures as clinical trial endpoints in ALS. Neurology 63, 1933–1935 (2004).

45 Kollewe, K. et al. ALSFRS-R score and its ratio: a useful predictor forALS-progression. J. Neurol. Sci. 275, 69–73 (2008).

46 Vender, R.L., Mauger, D., Walsh, S., Alam, S. & Simmons, Z. Respiratory systems abnormalities and clinical milestones for patients with amyotrophic lateral sclerosis with emphasis upon survival. Amyotroph. Lateral Scler. 8, 36–41 (2007).

47 Chiò, A., Hammond, E.R., Mora, G., Bonito, V., Filippini, G. Development and evaluation of a clinical staging system for amyotrophic lateral sclerosis. J Neurol Neurosurg Psychiatry. 86, 38–44 (2015).

48 Bozik, M.E., et al. A post hoc analysis of subgroup outcomes and creatinine in the phase III clinical trial (EMPOWER) of dexpramipexole in ALS. Amyotroph Lateral Scler Frontotemporal Degener. 15, 406–13 (2014).

49 Chen, X., et al. An exploratory study of serum creatinine levels in patients with amyotrophic lateral sclerosis. Neurol Sci. 35, 1591–7 (2014).

50 Van Eijk, R.P.A, Eijkemans, M.J.C., Ferguson, T.A., et al. Monitoring disease progression with plasma creatinine in amyotrophic lateral sclerosis clinical trials. J Neurol Neurosurg Psychiatry, 89:156–161 (2018).

